# Optimizing expression of Nanobody^®^ molecules in Pichia pastoris through co-expression of auxiliary proteins under methanol and methanol-free conditions

**DOI:** 10.1101/2023.03.31.535029

**Authors:** Manu De Groeve, Bram Laukens, Peter Schotte

## Abstract

**Background:** Ablynx NV, a subsidiary of Sanofi, has a long-standing focus on the development of Nanobody^®^ molecules as biopharmaceuticals (Nanobody^®^ is a registered trademark of Ablynx NV). Nanobody molecules are single variable domains, and they have been met with great success part due to their favorable expression properties in several microbial systems. Nevertheless, the search for the host of the future is an ongoing and challenging process. *Komagataella phaffi* (*Pichia pastoris*) is one of the most suitable organisms to produce Nanobody molecules. In addition, genetic engineering of Pichia is easy and an effective approach to improve titers.

**Results:** Here we report that *P. pastoris* engineered to co-express genes encoding four auxiliary proteins (HAC1, KAR2, PDI and RPP0), leads to a marked improvement in the expression of Nanobody molecules using the AOX1 methanol induction system. Titer improvement is mainly attributed to HAC1, and its beneficial effect was also observed in a methanol-free expression system.

**Conclusion:** Our findings are based on over a thousand fed-batch fermentations and offer a valuable guide to produce Nanobody molecules in *P. pastoris*. The presented differences in expressability between types of Nanobody molecules will be helpful for researchers to select both the type of Nanobody molecule and Pichia strain and may stimulate further the development of a more ecological methanol-free expression platform.

## Background

Single variable domains of heavy-chain only antibodies from Camelidae family members, known as Nanobody molecules, are particularly well-suited for the design of biopharmaceuticals [PMID: 33233943]. Compared to conventional antibodies, they can be easily engineered and cloned, allowing the generation of multi-specificity constructs. Furthermore, standard fed-batch fermentation using micro-organisms can express them and attain titers that are commercially viable. The *P. pastoris* expression platform is a well-established expression system capable of producing and secreting large quantities of Nanobody molecules. However, to improve cost of goods and the ecological footprint it is important to maximize the titer. To achieve this, many researchers have investigated ways to optimize the cultivation process and genetically engineer the host via co-expression of auxiliary proteins [PMID: 25080319, PMID: 29235217, PMID: 34037840, PMID: 29595107]. So far, no single auxiliary protein in *P. pastoris* has been identified that would be beneficial for every recombinant protein. Generally, multiple auxiliary proteins are usually screened and tested in combination with the recombinant protein of interest, or at best, with a similar type of proteins. This approach allows for a more comprehensive evaluation of which auxiliary proteins may be the most beneficial for the quality and titer of the recombinant protein. The complexity of determining whether an auxiliary protein is beneficial is further compounded by a multitude of other factors that must be taken into consideration such as the promoter used for both the auxiliary and recombinant protein, their copy number, their location in the genome, the secretion signal of the recombinant protein and many more [PMID: 22057543, PMID: 14755554, PMID: 11098467, PMID: 22885695; PMID: 11707618]. All these elements can significantly affect the efficacy of the auxiliary protein, making it difficult to accurately assess its beneficial properties. The Aox1 promoter has become a popular tool for driving the expression of recombinant proteins and subsequently many auxiliary proteins have been evaluated in this methanol-based system. Considering the safety risks associated with methanol exposure in a production plant, several groups have developed a methanol-free counterpart of the *P. pastoris* expression system [PMID: 28150747, PMID: 29280481, PMID: 30737736]. However, few of them investigated if and which auxiliary proteins can augment the expression levels of recombinant proteins in a methanol-free system [PMID: 27432633]. In this study, we compared the titer of a wide variety of Nanobody molecules expressed in wild type *P. pastoris* and in strains co-expressing genes encoding the auxiliary proteins in both methanol-based and methanol-free settings. After narrowing down the many proteins published to have a beneficial effect on expression, we chose four auxiliary proteins due to their distinct modes of action, i.e. HAC1, KAR2, PDI1 and RPP0 [PMID: 20591165, PMID: 16889384, PMID: 17051412, PMID: 18396911]. It is generally accepted that the rate of releasing recombinant molecules through the secretory route is influenced by folding dynamics. Therefore, our selection criteria for these four auxiliary proteins included their potential to reduce the intracellular accumulation of secreted proteins within the cell. *HAC1* and more specific the spliced *HAC1* mRNA encodes a transcription factor that activates transcription of genes involved in the unfolded protein response (UPR) [PMID: 34037840]. This response is activated when the cell is under stress from unfolded proteins in the endoplasmic reticulum (ER). The HAC1 upregulated UPR target genes encode a variety of potential auxiliary proteins such as chaperones, foldases, and proteins involved in glycosylation [PMID: 10847680, PMID: 34037840]. *KAR2* and *PDI1* are both UPR target genes which are upregulated by HAC1. They both enhance the expression of recombinant proteins by acting at different points in the folding process. KAR2 facilitates protein translocation and retention of unfolded or partially folded proteins, while PDI actively folds them and catalyzes the formation and reshuffling of disulfide bonds via its isomerase functions [PMID: 15990348]. The P0 protein, which is encoded by the acidic ribosomal protein RPP0, is essential for ribosome activity and is reported to increase secretion levels for multiple recombinant proteins. Even though RPP0 is not a transcription factor, its overexpression can change the levels of many proteins, including chaperones [PMID: 15866509, PMID: 11923285, PMID: 18396911, PMID: 8195220]. We found that Nanobody titers are generally improved when they are co-expressed with one or more of these auxiliary proteins in both the methanol-based and methanol-free expression systems. The results of this study will likely enable biopharmaceutical companies to develop cost-effective production processes for Nanobody molecules and facilitate the clinical development of these promising molecules.

## Results

### *Assessing the individual effect of* RPP0, KAR2, PDI1 and HAC1 *on titer of the low expressing Nanobody molecule A using the methanol-based expression system*

Nanobody molecule A is a bivalent Nanobody molecule consisting of two variable domains from two heavy-chain llama antibodies. The N-terminal building block of the molecule is a VHH1 type immunoglobulin single variable domain which typically contains 2 disulfide bridges. While the C-terminal subunit is a VHH3 type immunoglobulin single variable domain and contains one disulfide bridge. The subunits are fused head-to-tail with a 9GS linker. In a generic fed-batch fermentation and under the control of the Aox1 promoter the expression level of Nanobody molecule A in wild type *P. pastoris* X-33 strain is low (0.1 g |^-1^).

For this reason, we explored whether the titer could be improved when auxiliary proteins such as RPP0, KAR2, PDI1 and HAC1 were co-expressed. Therefore, we super-transformed the *P. pastoris* clone expressing the Nanobody molecule A with these individual auxiliary proteins. A clone from each transformation was selected and evaluated in fed-batch fermentation. Titers were 3-, 5-, 14- and 15-fold higher than the reference clone when respectively RPP0, KAR2, PDI1 and HAC1 were co-expressed (Table 1). Based on these results we decided to make a *P. pastoris* strain containing these four auxiliary proteins that could be used as a superior platform strain for the expression of other Nanobody molecules.

**Table 1.** Evaluation of HAC1, PDI1, RPP0 and KAR2 for titer improvement of Nanobody molecule A.

**Evaluation of HAC1, PDI1, RPP0 and KAR2 for titer improvement of** 677 **Nanobody molecule A.**
| Strain | Titer (g l<sup>-1</sup>) |
| --- | --- |
| X-33 | 0,1 |
| X-33 + RPP0 | 0,3 |
| X-33 + KAR2 | 0,5 |
| X-33 + PDI1 | 1,4 |
| X-33 + HAC1 | 1,5 |

Productions were performed at 2 l scale, pH 6, 30 °C in complex medium with a
methanol feed rate of 4 ml l<sup>-1</sup> h<sup>-1</sup>. Indicated titers are measured in clarified cell broth via SDS-PAGE/Coomassie staining and quantification of bandvolume measured using ImageQuant software (Cytiva).

Because the X-33 expression system is a commercial system and comes with a R&D license cost we decided to generate a strain containing the four auxiliary proteins in the NRRL Y-11430 background which was acquired from the United States Department of Agriculture (USDA-ARS).

### *Improved expressability of Nanobody molecules in the strain* NRRL Y-11430 co-expressing *RPP0, KAR2, PDI1 and HAC1 using the methanol-based expression system*

To generate the NRRL Y-11430 strain co-expressing genes of the four auxiliary proteins we used standard cloning techniques as detailed in the Materials and Methods section. A clone was selected that demonstrated the presence of the four auxiliary proteins via PCR. To evaluate if the new clone showed improved expression capabilities versus the wild type parental strain we tested the titers of two Nanobody molecules B and C, in both strains under the control of the Aox1 promoter. Nanobody molecule B is a trivalent Nanobody molecule and the building blocks are linked together via 35GS linkers. Nanobody molecule C is a tetravalent Nanobody molecule, consisting of four Nanobody building blocks linked together with 5GS-9GS-9GS linkers. To evaluate if co-expression of auxiliary proteins would be affected by the type of secretion signal we cloned Nanobody molecules B and C in frame with respectively the α-mating factor secretion signal (aMF) from S. cerevisiae secretion [PMID: 6087338] or the secretion signal from the cell wall protein CWP1 [PMID: 27287536] (GeneID:8199470). The expression vectors were transformed to the wild type NRRL Y-11430 strain and the strain co-expressing genes encoding for the four auxiliary proteins. Clones were randomly picked from zeocin plates and tested in fed batch fermentation. We observed a significant improvement in titer for both Nanobody molecules in the NRRL Y-11430 strain co-expressing *RPP0, KAR2, PDI1* and *HAC1* (Table 2), thus providing confidence that the new strain is superior to the wild type NRRL Y-11430 strain in terms of Nanobody molecule expression. Based on these results we decided to switch from the wild type *P. pastoris* strain to the strain expressing the four auxiliary proteins for generic production of our Nanobody molecules. Data on titer from more than 1000 different Nanobody molecules produced in fed-batch fermentation was collected over a period of several years and presented in Table 3. This is a remarkable data set because it is the first time that this much information on titers using fed-batch fermentation mediated production of Nanobody molecules is presented. In addition, fermentation data is more useful for pharmaceutical development than the titers obtained from shake flasks or small scale 96-deepwell expression. These methods of assessing expressability are not very reliable when predicting the titers that can be achieved in fed-batch fermentation between clones and the expressability of Nanobody formats. Furthermore, our data demonstrate that Nanobody molecules generally express well in wild type *P. pastoris* with an average titer of 1.4 g |^-1^ in NRRL Y-11430 (N = 54). Interestingly, the titer of Nanobody molecules expressed in the wild type *P. pastoris* strain tend to drop when the format of the Nanobody molecule contains more than 2 building blocks. This trend is not observed with the strain co-expressing the four auxiliary proteins. Although based on a limited set of 40 different Nanobody molecules, it is noteworthy that the average titer of Nanobody formats containing a VHH1 building block expressed in the wild type X-33 *P. pastoris* strain is significantly (p=0.0087) lower than non-VHH1 formats (1.1 g |^-1^ versus 1.7 g |^-1^) (Table 4). However, like higher valency Nanobody molecules, the average titer of VHH1 containing formats can be increased to 2.7 g |^-1^ (N = 150) when co-expressing *RPP0, KAR2, PDI1* and *HAC1* (Table 4). In addition, the difference in average titer of VHH1 and non-VHH1 formats is much smaller and not statistically significant (p=0.0916) in this improved Pichia strain (2.7 g |^-1^ versus 3.1 g |^-1^). We can conclude that the strain co-expressing the genes encoding the four auxiliary proteins in a methanol-based expression system is a suitable production host for a wide range of Nanobody molecules, as it is capable to reach high titers for multi-valent Nanobody molecules compared to the wild type *P. pastoris* strain.

**Table 2.**
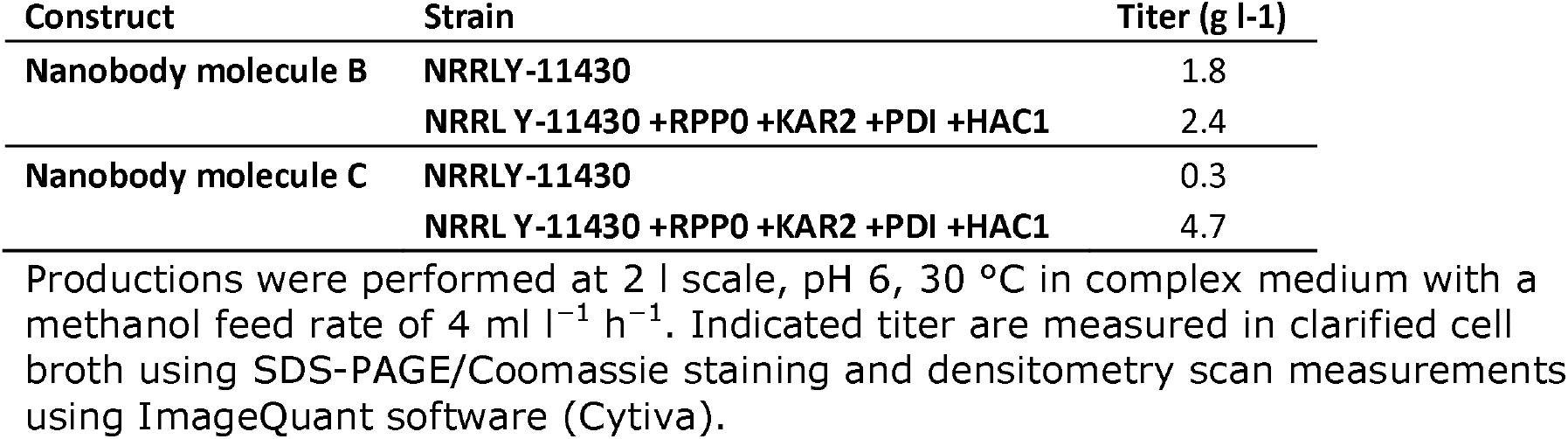
Evaluation of the base strain NRRL Y-11430 co-expressing *RPP0 + KAR2 + PDI + HAC1* for titer improvement of Nanobody molecule B and C.

| Construct | Strain | Titer (g l<sup>-1</sup>) |
| --- | --- | --- |
| Nanobody molecule B | NRRL Y-11430 | 1.8 |
|  | NRRL Y-11430 +RPP0 +KAR2 +PDI +HAC1 | 2.4 |
| Nanobody molecule C | NRRL Y-11430 | 0.3 |
|  | NRRL Y-11430 +RPP0 +KAR2 +PDI +HAC1 | 4.7 |

Productions were performed at 2 l scale, pH 6, 30 °C in complex medium with a methanol feed rate of 4 ml l<sup>-1</sup> h<sup>-1</sup>. Indicated titer are measured in clarified cell broth using SDS-PAGE/Coomassie staining and densitometry scan measurements using ImageQuant software (Cytiva).

**Table 3.** Titer of different Nanobody molecules according to their format and expressed in different strains.

| System | Strain | Average titer (g l<sup>-1</sup>) | Average titer (g l<sup>-1</sup>) |  |  |  |  |  |
| --- | --- | --- | --- | --- | --- | --- | --- | --- |
|  |  |  | Nanobody valency |  |  |  |  |  |
|  |  |  | mono | bi | tri | tetra | penta | hexa |
| Methanol-based | NRRL Y-11430 | 1.4<br>(N=54) | 2.7<br>(N=13) | 1.8<br>(N=4) | 0.8<br>(N=18) | 1.1<br>(N=17) | 0.8<br>(N=2) | -<sup>a</sup> |
|  | NRRL Y-11430 +RPP0 +KAR2 +PDI +HAC1 | 3.1<br>(N=849) | 2.7<br>(N=314) | 3.2<br>(N=82) | 3.2<br>(N=148) | 3.4<br>(N=172) | 3<br>(N=102) | 4.1<br>(N=31) |
| Methanol-free | NRRL Y-11430 | 0.6<br>(N=16) | - | 0.4<br>(N=2) | - | 0.5<br>(N=11) | 1.4<br>(N=3) | - |
|  | NRRL Y-11430 + HAC1 | 3.7<br>(N=156) | 3.4<br>(N=93) | 4.3<br>(N=12) | 4.5<br>(N=12) | 3.7<br>(N=30) | 4.6<br>(N=8) | 3.6<br>(N=1) |

<sup>a</sup> - = no data available
Productions were performed under generic fed-batch fermentations as described in the material and methods. Indicated titer are measured in clarified cell broth. Nanobody valency reflects the number of genetically fused Nanobody building blocks in the construct.

**Table 4.**
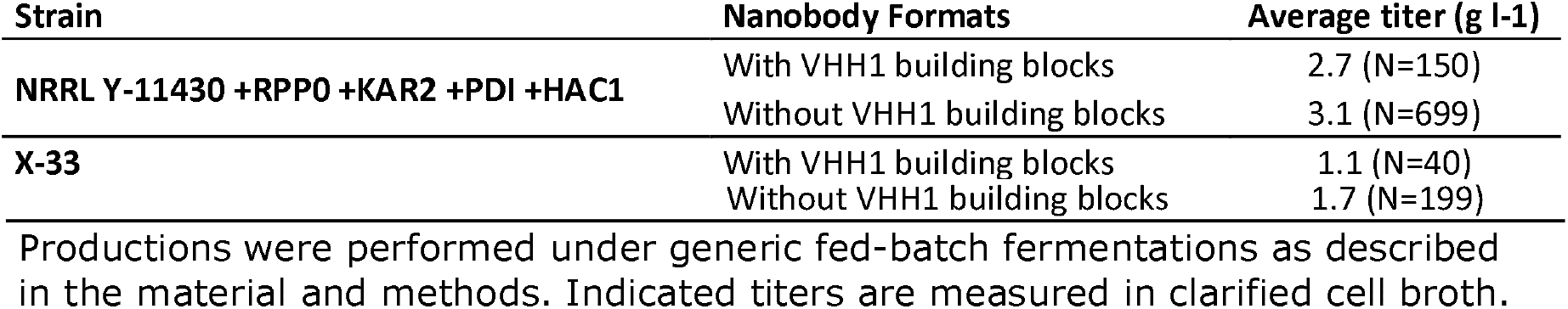
Titer of Nanobody molecules constructed with or without VHH1 building blocks in their format.

### *Improved expressability of Nanobody molecules in the strain* NRRL Y-11430 *co-expressing HAC1 using methanol-free expression system*

In recent years we observe an increasing interest in methanol-free *P. pastoris* expression systems due to the operational advantages they offer over the widely used AOX1 methanol-based system. The primary benefit is the improved safety conditions that result from the elimination of methanol, which is volatile, flammable, and toxic. Methanol-free production plants have a simpler design avoiding ATEX proof areas for methanol and are therefore more cost-effective, making them an attractive option for pharmaceutical companies and contract manufacturers. Currently, there are several methanol-free expression systems available for *P. pastoris* [PMID: 23165100]. Depending on the recombinant protein, several of these methanol-free promoters showed equal or even better expression levels than the strong Aox1 promoter. Here we evaluated the commercially available methanol-free promotors of Bisy which include the carbon source regulated promoters Pdc and Pdf [PMID: 30735181, PMID: 32100120]. We evaluated the titers of Nanobody molecules in this system and determined if co-expression of genes coding for auxiliary proteins could enhance the titer. We focused on the effect of co-expressing *HAC1* because it showed the highest improvement in expression of Nanobody molecule A in our first experiments. A base strain was generated containing the *Pac-HAC1* expression cassette in the NRRL Y-11430 strain. Subsequently, the wild-type NRRLY-11430 and *HAC1*-containing NRRL Y-11430 strains were transformed with expression vectors coding for five different Nanobody molecules (D, E, F, G and H) under the control of the Pdf promoter. All five Nanobody molecules were multivalent formats containing 4 to 6 Nanobody building blocks separated with GS linkers. Expression was tested in fed-batch fermentation according to our generic methanol-free cultivation conditions as described in the materials and methods. Titer of all five Nanobody molecules ranged between 0.5 and 1.6 g |^-1^ cell broth (Figure 1). For all 5 Nanobody molecules we observed a significant increase in titer when *HAC1* is co-expressed reaching titers up to 4 g |^-1^ cell broth. The data clearly show the beneficial effect of HAC1 in the methanol-free expression system.

**Fig. 1.**
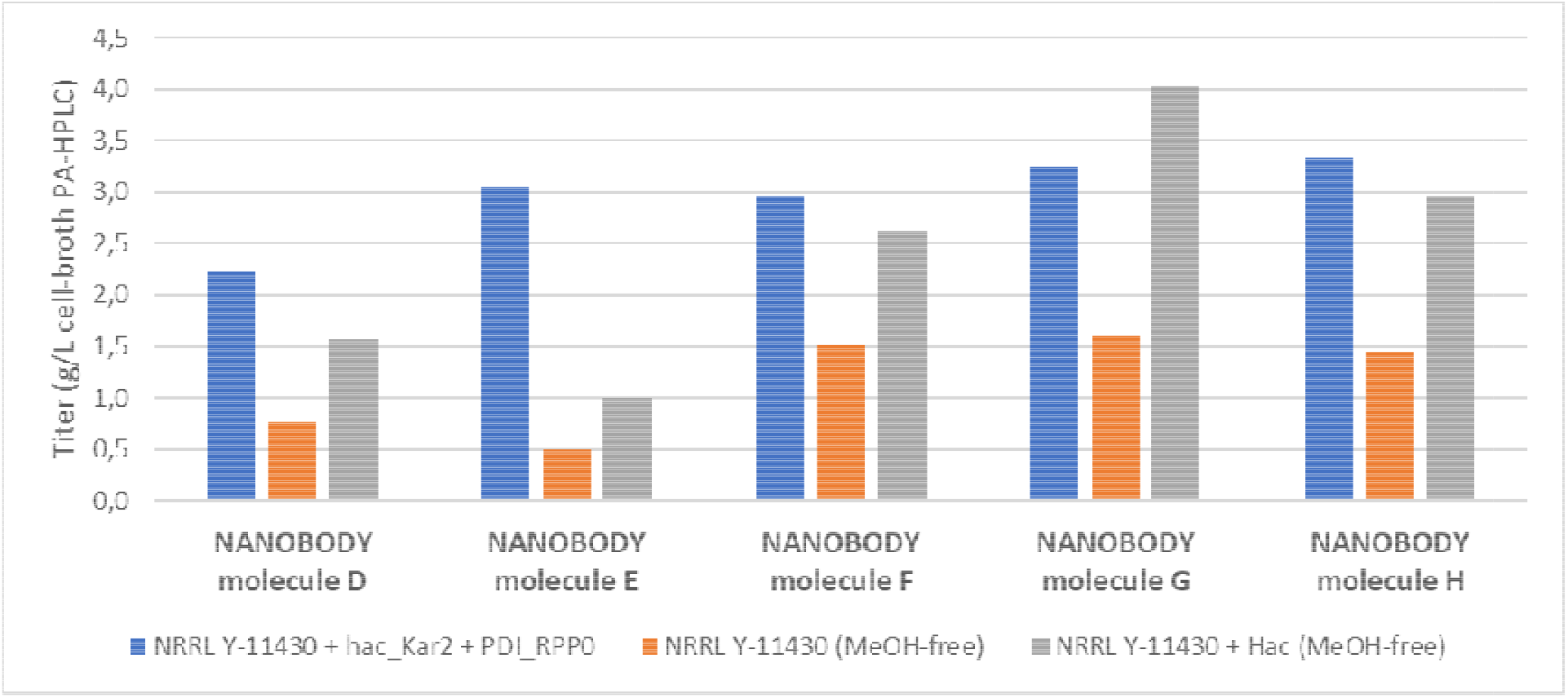
5 Nanobody molecules (D, E, F, G and H) were used to benchmark the methanol-free platform strain co-expressing *HAC1*. As a reference the 5 Nanobodies were expressed under methanol-free conditions in the wild type *P. pastoris* and in the methanol-based system co-expressing *RPP0, KAR2, PDI1* and *HAC1*. Cultivation were performed in fed-batch fermentation according to a generic methanol-based or methanol-free cultivation conditions as described in materials and methods. Indicated titers in clarified cell broth are measured using PA-HPLC.

As a reference we also expressed the 5 Nanobodies in the methanol-based system using the NRRL Y-11430 co-expressing *RPP0, KAR2, PDI1* and *HAC1*. 3 Nanobody molecules reached a similar titer in both systems and for 2 molecules the methanol-based system was superior. Taking the benefits of methanol-free expression into account, we decided once again to switch our generic methanol-dependent host in favor of the methanol-free *HAC1*-containing NRRL Y-11430 strain. Thus far we have evaluated the titers in fed-batch fermentation of 156 different Nanobody molecules in this new strain (Table 3). The data indicate that generally the *HAC1* co-expressing strain is superior to the wild type strain with average titer of respectively 3.7 g |^-1^ versus 0.6 g |^-1^. In addition, the data also indicate that the average titer of different Nanobody formats in the methanol-free system surpasses that of the methanol-based system with 4 auxiliary proteins (3.7 g |^-1^ vs 3.1 g |^-1^). Interestingly, the titers of Nanobody molecules remain relatively stable in relation to their size in the methanol-free *HAC1-* containing NRRL Y-11430 strain similar to what we observe for the methanol-based strain expressing the four auxiliary proteins.

## Discussion

30 Years after the serendipitous discovery of Nanobody molecules [PMID: 8502296] their popularity has grown exponentially. Main factor is the ability to genetically fuse Nanobody building blocks and as such within a single design generate multiple antigen recognition. Their ease of production using microbial expression systems, as demonstrated in this study, has further attracted the attention of several pharmaceutical companies.

The structure of a Nanobody building block is simple and may have good folding kinetics, however, this may not be the case for larger multivalent Nanobody formats. Our data indicate that the folding and secretion machinery of wild type *P. pastoris* can handle bivalent Nanobody molecules without a significant decrease in titer compared to monovalent building blocks. However as of trivalent Nanobody molecules formats a decrease in titer can be observed. This could be linked to the process of disulfide bridge formation which for larger Nanobody formats may become increasingly complex and could lead to slower folding kinetics, protein aggregation and low titers. This idea is supported by our data that Nanobody formats which include VHH1 building blocks, having 2 disulfide bridges in their structure, are expressed at lower titers. Several groups report that a more oxidative state of the ER could be beneficial for protein folding and titer improvement [PMID: 22406321, PMID: 28357216]. However, having a good titer for a Nanobody molecule is likely to involve additional factors because they can be folded and secreted without proper oxidized disulfide bridges as well. Depending on the Nanobody molecule we typically detect 5 to 20% of the secreted molecules lacking one or more disulfide bridges (unpublished data). Incorrect protein folding can cause ER stress and an unfolded protein response (UPR), which leads to the induction of stress genes such as auxiliary proteins [PMID: 22116877]. In a wild type *P. pastoris* strain, the UPR response may not be adequate to deal with a large amount of difficult to express Nanobody molecules. Perhaps somewhat surprisingly, but luckily, the UPR response itself can be boosted or mimicked by overexpressing UPR regulated genes under the control of a similar strong and inducible promoter system as used for the Nanobody molecules. One may expect that overexpressing of genes that are critical in protein expression could dysregulate the entire machinery leading to worse titers and such observations have been described [PMID: 32175802]. In our first assessment co-expressing genes coding for four auxiliary proteins with Nanobody molecule A we did not observe such effect and therefore decided to go ahead in making a platform strain containing the four auxiliary proteins. Data from 699 fed-batch fermentations indicate that this platform strain improved titers by an average of 100%, resulting in an average titer of 3.1 g |^-1^. However, 6% of Nanobody molecules still had low titers (below 0.5 g |^-1^) regardless of their format (data not shown). A similar observation can be made for the methanol-free HAC1 base strain in which 4% of Nanobody molecules had low titers. However, we cannot exclude that the titers of these low expressing Nanobody molecules are still better than if they would have been expressed in the wild type *P. pastoris* strain. Nevertheless, it remains remarkable that for such similar molecules there is a broad spread in titer ranging from 0.1 g |^-1^ to more than 10 g |^-1^. This may indicate that other more specific auxiliary proteins would be needed for these difficult to express Nanobody molecules. For the expression of Nanobody molecule A the individual contribution of RPP0 and KAR2 was significantly lower than for PDI1 and HAC1. We can only speculate that the improved expression of Nanobody molecule A by co-expressing *PDI1* is due to its isomerase activity which actively participates in the folding of the 3 disulfide bridges of that molecule. So far it remains to be proven that during the Nanobody folding process the free sulfhydryl groups interact with *P. pastoris* foldases and isomerases such as PDI. It is somewhat remarkable that overexpression of PDI as one of the most abundant and stable proteins in the ER still exerts effect on titer [PMID: 7287643, PMID: 1988050]. Some report that overexpression of PDI may lead to the induction of an unfolded protein response (UPR) [PMID: 17051412]. The fact that HAC1 as a transcription factor increased the titer of Nanobody molecule A slightly better than PDI1 could be related to HAC1 mediated upregulation of endogenous PDI1 and additional UPR genes. Due to the distinct biology and metabolism of cells grown on either methanol or glycerol, we did not assume that the same auxiliary proteins would be equally effective in facilitating expression of Nanobody molecules in a methanol-based and a methanol-free system. Nevertheless, we decided to take a risk and evaluate only HAC1 in the methanol-free system, as it may have a pleiotropic effect. The expression profile of the five Nanobody molecules D, E, F, G and H was compared in both the methanol-free system co-expressing *HAC1*, and the methanol-based system co-expressing the four auxiliary genes. These experiments confirmed that the titer for the same Nanobody molecule can be different depending on the promoter system and the co-expressed auxiliary proteins. Nevertheless, with an average titer of 3.7 g |^-1^ for different Nanobody molecules the methanol-free base strain co-expressing *HAC1* has an attractive feature. In fact, we occasionally observed high titers reaching double digits with this strain, even for penta- and hexavalent Nanobody molecules.

For the ease of working our Nanobody expression platform standardly uses the aMF secretion signal. The aMF contains a pre-peptide that targets proteins into the endoplasmic reticulum and a pro-peptide which regulates efficient secretion [PMID: 27984193]. Occasionally we observe inefficient maturation of the aMF which may result in product heterogeneity for a clinical lead Nanobody molecule. Removing the aMF pro-sequence or using an alternative secretion signal can sometimes solve this problem [PMID: 23160737, PMID: 31202796]. However, and for unknown reasons, we tend to see a drop in titer when using alternatives secretion signals (unpublished data). The expression data of Nanobody molecule C shows that co-expression of the tested auxiliary proteins can improve the titer when using alternative secretion signals such as CWP1 secretion signal in both the methanol and methanol-free expression system.

For the development of a biopharmaceutical it is important to have an efficient production process including a high titer. Firstly, high titers ensures that sufficient drug substance can be produced in one production campaign reducing development timelines and facilitating a speedy approval process, leading to quicker access to the product for patients. Secondly, it reduces the cost of goods as more product can be produced in a shorter period. Furthermore, there are environmental benefits from an optimal production process, as fewer energy, water and chemicals need to be used, resulting in fewer emissions and a smaller environmental footprint.

## Conclusion

This research aimed to summarize our observations from over a decade of strain testing for the expression of Nanobody molecules in *P. pastoris*, and provides useful findings for future research in making Nanobody based biopharmaceuticals. The protein secretion machinery of *P. pastoris* is comparatively less efficient than that of mammalian cells [PMID: 24390926], However, our data suggests that Nanobody molecules and *P. pastoris* are well-suited to each other. We anticipate that further research into the molecular mechanisms of how and why auxiliary proteins contribute to increased Nanobody expression will lead to big improvements in this area.

## Methods

### Generation of strains expressing Nanobody molecule A

Top10 cells were used for the amplification of recombinant plasmid DNA (Invitrogen Corp., Carlsbad, CA). The coding information of Nanobody molecule A was cloned into the multiple cloning site of a *P. pastoris* expression vector (derivate of pPIC6a, Invitrogen) that contains a blasticidin resistance marker, such that the Nanobody sequence was downstream of and in frame with the aMF signal peptide sequence. To generate *P. pastoris* clones with more than 1 copy number of the expression cassette in the genome, a unique BglII site in the *P. pastoris* expression vector was used to introduce a second expression cassette of Nanobody molecule A. The expression vectors used for the co-expression of *HAC1, KAR2, PDI1* and *RPP0* with Nanobody molecule A carried the zeocin selection marker. The expression vectors of all other Nanobody molecules tested in this paper were cloned with a single expression cassette and used zeocin as a selection marker. Transformation was performed by standard techniques as previously described [PMID: 27267127].

### Generation of the NRRL Y-11430 base strains containing auxiliary proteins

For the construction of the *4 auxiliary proteins* base strain in a NRRL Y-11430 background we used the pPpT4AlphaS vector backbone [PMID: 22768112]. The NRRL Y-11430 strain was retrieved from the United States Department of Agriculture (USDA) and Agricultural Research Service (ARS). The coding sequence of the spliced *HAC1* gene was ordered as 2 pieces of synthetic gene fragments (gblocks) at Integrated DNA Technologies, Inc.. Assembly of the 2 synthetic gene fragments was done by PCR and the *HAC1* coding sequence was cloned into the pPpT4AlphaS expression vector under the control of the Aox1 promoter. This expression vector has the plasmid reference ID number P-16.76. The recognition site for the restriction enzyme BsiWI was removed in plasmid P-16.76 resulting in a new plasmid reference ID number P-18.46. The coding sequence of the *KAR2* gene (GenBank: SOP81505.1) was isolated from the wild type strain NRRL Y-11430 strain by PCR and cloned into an expression vector under the control of the Aox1 promoter (plasmid reference ID number P-16.77). The construct P-16.77 was used to isolate the Kar2 expression cassette by PCR which was subsequently cloned into the plasmid P-18.46. This resulted in the final construct P-18.48 containing the genes for both *HAC1* and *KAR2* each under control of the Aox1 promoter. The vector contains a blasticidin selection marker which enabled the selection of a *P. pastoris* clone expressing the *HAC1* and the *KAR2* genes. The coding sequence of the *RPP0* gene (GenBank: CCA36233.1) was isolated from the wild type strain NRRL Y-11430 strain by PCR and cloned under the control of the Aox1 promoter (plasmid reference ID number P-16.79). The recognition site for the restriction enzyme BsiWI was removed in plasmid P-16.79 resulting in a new plasmid reference ID number P-18.47. The coding sequence of the *PDI1* gene (GenBank: CCA40283.1) was isolated from the wild type strain NRRL Y-11430 strain by PCR and cloned into an expression vector under the control of the Aox1 promoter (plasmid reference ID number P-16.78). Via overlap PCR the BamHI site in the *PDI1* gene was removed prior to cloning into the plasmid P-18.47. This resulted in the final construct P-18.50 containing the RPP0 and *PDI* genes each under control of the Aox1 promoter. The vector contains a Blasticidin selection marker which enabled for the selection of a *P. pastoris* clone expressing *RPP0* and *PDI1*. Both constructs P-18.50 (PDI + RPP0) and P-18.48 (HAC1 + KAR2) were stably co-transformed into the genome of the NRRL Y-11430 strain generating the strain NRRL Y-11430 + RPP0 + KAR2 + PDI + HAC1 which is subsequently used for the transformation and expression of Nanobody molecules.

Generation of the methanol-free base strain NRRL Y-11430 + HAC1 was performed as follows. Starting from the earlier described vector P-16.76 the Aox1 promoter was replaced by the Pdc promoter [PMID: 26592304] from the P. pastoris catalase *CTA1* gene ordered as a clonal gene plasmid at Twist (Twist Bioscience). Cloning of the expression vector was performed using NEBuilder HiFi DNA Assembly Master Mix (Gibson cloning) and was performed according to manufacturer’s guidelines (cat. n° E2621L, New England Biolabs). In addition, the Blasticidine selection marker was replaced by a marker conferring resistance to the Geneticine for selection. The final plasmid reference had ID number P25.57. Following linearization of plasmid P-25.57, the construct was stably transformed into the genome of the NRRL Y-11430 strain generating the basic strain NRRL Y-11430 + HAC1 which is subsequently used for the transformation and expression of Nanobody molecules under methanol-free conditions. The expression vector used for the expression of Nanobody molecules under methanol free conditions was made by exchanging the Aox1 promotor in the P-16.76 vector with the *H. polymorpha* Pdf promoter [PMID: 26592304] ordered at Twist (Twist Bioscience) and exchanging the Blasticidine selection marker to Zeocin. As such both Pdc- and Pdf-promoters are derepressible promoters, controlling expression of the *HAC1* gene and the Nanobody molecule in a MeOH-free process.

### Generic fed-batch fermentation conditions

High cell density fed-batch fermentation for production of Nanobody molecules was performed at 0.25 L (Ambr250 MO units, Sartorius Stedium Biotech), 2 L or 5 L scale (Biostat B-plus and B-DCU units, Sartorius Stedium Biotech) as previously described [PMID: 9680637].

During expression of Nanobody molecules, the temperature in the bioreactor was controlled at 30°C, dissolved oxygen at 30% and pH at 6.0.

For methanol-based processes, expression of the Nanobody molecules was induced by addition of 100% methanol to the bioreactor at a constant feeding rate of 4 mL/h/L initial volume for approximately 96 hours. For methanol-free processes, expression of the Nanobody molecules was derepressed by addition of a 80% (w/w) glycerol feed for 80-96 hours at a limiting and decreasing feeding rate (at start of derepression phase: 15 g/h/L initial volume, 4.5 h after start: 8 g/h/L, 9 h after start: 4 g/h/L, 62 h after start until end of fermentation: 2 g/h/L).

### Titer determination

At end of fermentation, cell broth samples were centrifuged (11285 g; 20 minutes) and titers in the cell-free supernatant were quantified using SDS-PAGE/Coomassie staining and densitometry scan measurements using ImageQuant software (Cytiva). For Nanobody molecules C, D, E, F and G in Figure 1 titer was determined using a PA-HPLC analysis using a POROS A 20 μm column (Applied Biosystems) on an Agilent 1260 system. Briefly, clarified broth sample was centrifuged to remove any remaining particulate matter. The sample was applied on the column according to the manufacturer’s instructions. The following buffers/solutions were used: mobile phase A (0.1 M Phosphate, 0.15 M NaCl, pH7), mobile phase B (0.5 M NaCl concentration and a pH of 1.9). In short, mobile phase A is used to create optimal binding conditions on the column. Mobile phase B is used to elute the Nanobody molecule from the column. The dilution buffer, which is used for blank injections and to dilute samples, is made from mobile phase A and 0.05% (w/v) Tween 20. The column compartment temperature was set at 20°C. When the samples are eluted from the column they were detected with UV-DAD detector with the standard setting of 280 nm used.

### Chemicals

Enzymes were purchased from New England Biolabs.

Difco^™^ yeast nitrogen base without amino acids (YNB), HypA peptone was obtained from BioSpringer and yeast extract from Oxoid.

Glucose was obtained from Merck chemicals, glycerol from VWR chemicals, sorbitol and D-Biotin from Sigma Aldrich. Select agar and Zeocin™ were obtained from Invitrogen. Blasticidine was purchased from InvivoGen (Cayla InvivoGen Europe). Geneticin was purchased from Thermo Fisher Scientific Inc..

### Statistical analysis

The non-parametric significance of the differences between VHH1 and non-VHH1 populations was determined according to the Wilcoxon rank test in JMP 16.2 software.

## Data availability

All data are included in the manuscript. Further queries or additional information can be requested to the corresponding author.

## Abbreviations

aMF: α-Mating factor secretion signal
Aox: Alcohol oxidase
ATEx: Atmosphere Explosible
HAC: Homologous to ATF and CREB
CWP: Cell wall protein
ER: Endoplasmic reticulum
GS: Glycine-Serine
KAR: Karyogamy
OD: Optical density
PA-HPLC: protein A high-performance liquid chromatography
PDI: Protein disulfide isomerase
PCR: Polymerase chain reaction
PDC: De-repressible catalase promoter
PDF: De-repressible FMD promoter from *Hansenula polymorpha*
RPP0: Ribosomal protein P0
SDS-PAGE: Sodium dodecyl-sulfate polyacrylamide gel electrophoresis
UPR: Unfolded protein response
VHH: Single variable domain on a heavy chain

## Author information

All authors are employed at Sanofi Ghent, Ablynx NV, Technologiepark 21, 9052, Zwijnaarde, Belgium

## Authors’ contributions

MDG, BL and PS performed data mining and designed and interpreted the data. PS wrote the manuscript. All authors read and approved the final manuscript.

## Acknowledgements

We thank Yannick Van Haelst for assistance in statistical analysis. The authors are grateful to the following individuals for reviewing and their contributions to this work: Pauline Dechaene, Katrien Vlassak, Ann Brigé and Rebecca Sendak.

